# Hayai-Annotation Plants: an ultra-fast and comprehensive gene annotation system in plants

**DOI:** 10.1101/473488

**Authors:** Andrea Ghelfi, Kenta Shirasawa, Hideki Hirakawa, Sachiko Isobe

## Abstract

Hayai-Annotation Plants is a browser-based interface for an ultra-fast and accurate gene annotation system for plant species using R. The pipeline combines the sequence-similarity searches, using USEARCH against UniProtKB (taxonomy Embryophyta), with a functional annotation step. Hayai-Annotation Plants provides five layers of annotation: 1) gene name; 2) gene ontology terms consisting of its three main domains (Biological Process, Molecular Function, and Cellular Component); 3) enzyme commission number; 4) protein existence level; 5) and evidence type. In regard to speed and accuracy, Hayai-Annotation Plants annotated *Arabidopsis thaliana* (Araport11, representative peptide sequences) within five minutes with an accuracy of 96.4 %.

**Availability and Implementation:** The software is implemented in R and runs on Macintosh and Linux systems. It is freely available at https://github.com/kdri-genomics/Hayai-Annotation-Plants under the GPLv3 license.

## 1. Introduction

The main goal in plant science is to understand plant biological systems in order to describe patterns of evolution and diversity. Because genome information would accelerate the process of plant research, it is crucial that molecular biologists have broad and accurate knowledge of gene profiles in the relevant genomes. There are many tools available for gene and protein function annotation. To cite a few, Blast2GO (Conesa and Götz, 2008), TRAPID (Van Bel, et al., 2013), Mercator 4 (Lohse, et al., 2014), and FunctionAnnotator (Chen, et al., 2017). They use different algorithms for sequence alignments, respectively, BLAST (Altschul, et al., 1997), RAPSearch (Ye, et al., 2011), MapMan (Thimm, et al., 2004) and LAST (Frith, et al., 2010). However, they require a long time to run and the annotation assignment does not consider the type of evidence that supports the existence of a protein (Protein Existence Level). In UniProtKB (UniProt Consortium, 2018) there is five types of evidence for the existence of a protein: 1. experimental evidence at a protein level, 2. experimental evidence at a transcript level, 3. protein inferred by homology, 4. protein predicted, 5. protein uncertain. To address these issues, we developed Hayai-Annotation Plants, which is an ultra-fast, highly accurate and comprehensive gene annotation system in plants.

Hayai-Annotation Plants is based on sequence similarity searches using USEARCH (Edgar, 2010), which is an algorithm orders of magnitude faster than BLAST, against UniProtKB, taxonomy Embryophyta (land plants). Hayai-Annotation Plants makes use of the complete set of protein information from UniProtKB to provide five layers of annotation: gene name; Gene Ontology (Ashburner, et al., 2000) (GO) consisting of three main categories (Biological Process, Molecular Function and Cellular Component); Enzyme Commission (EC) number; protein existence level; and evidence type. Hayai-Annotation Plants introduces an algorithm that gives priority for higher levels of protein existence, followed by curated evidence types, rather than evidences inferred from electronic annotation (evidence types describe the source of the information in UniProtKB). We believe this algorithm has the potential to standardize and increase GO and EC annotation assignments when reliable database is used.

## 2. Material and Methods

Hayai-Annotation Plants is an R package that employs the R-package “Shiny” for its browser interface, as well as the free version of USEARCH (32-bits) (Edgar, 2010) for sequence alignment, and uses UniProtKB (taxonomy Embryophyta) to designate the gene name, GO term and code, EC number, protein existence level, and evidence type. FASTA-formatted file is required as minimum input, and five parameters can be customized (type of alignment – local or global, maximum hits per query, minimum sequence identity, minimum query coverage, and E-value). In addition, we developed two types of algorithms to assign annotation. The “Alignment Score” algorithm gives priority to sequence identity parameter, followed by protein existence, then evidence type. The “Protein Existence Level” algorithm has another priority order, first consider the highest protein existence level parameter, followed by evidence type, then sequence identity (for global alignment) or score (for local alignment).

The annotation process can be separated in two steps, the first is the sequence alignment and the second the functional annotation step, with the incorporation of gene name, GO terms and codes, EC number, existence level, and evidence type. In the first step we implemented USEARCH, but since the free version of USEARCH only allows files smaller than 4 GB, we split the database into four uniform parts (same number of sequences each). In the functional annotation step, each query with its UniProt code associated is merged with the information from UniProtKB, following the priority given by the selected type of algorithm. Figure 1 shows a schematic description of the pipeline. Since not all subjects have a complete set of annotation, and in order to increase the level (number of layers) of annotation, within the parameters chosen and depending on the selected algorithms, Hayai-Annotation Plants will select GO information, for each main category, independently. Thus, if one subject does not have information regarding a particular GO category (BP, MF or CC) it will look for the next closest subject automatically until it finds an entry that holds the required information. This means that one query can use more than one UniProt code to assign annotation.

**Figure 1.**
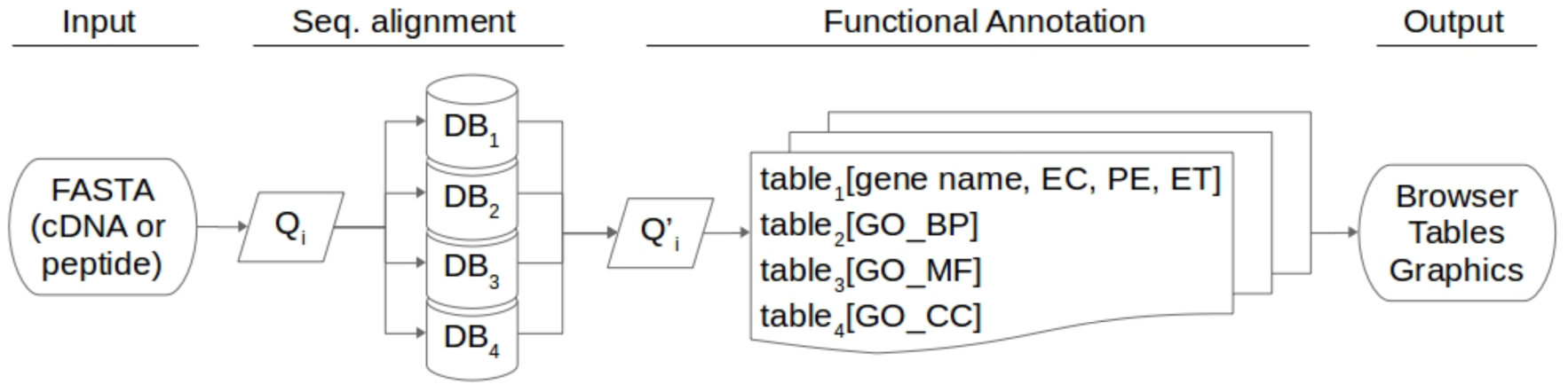
Hayai-Annotation Plants pipeline. Q: query sequence; index i: each query sequence; DB_1_, DB_2_, DB_3_, DB_4_: quarters of UniProtKB (taxonomy Embryophyta); EC: Enzyme Commission; PE: Protein Existence; ET: Evidence Type; GO_BP, GO_MF, GO_CC: Gene Ontology tables, containing information of Biological Process, Molecular Function and Cellular Component, respectively.

Hayai-Annotation generates six tables and four graphics, that can be easily download all at once using a “Download” link at the browser interface. The main table created by Hayai-Annotation Plants selects the unique functional annotation for each query sequence. Regarding GO terms for each main domain (BP, MF, and CC) and EC number, Hayai-Annotation Plants provides graphics with the top 50 terms/number and tables of all results. Besides, Hayai-Annotation Plants makes another file that can be directly uploaded to KEGG Mapper (Du, et al., 2014) (see https://github.com/kdri-genomics/Hayai-Annotation-Plants for more details).

## 3. Results and Discussion

To compare speed and accuracy of the annotation tools, we annotated the genes predicted in *Arabidopsis thaliana* genome (Araport11, representative peptide sequences) using Hayai-Annotation Plants, Blast2GO, TRAPID, Mercator 4, and FunctionAnnotator, and calculate Annotation Edit Distance (AED) based on the manually curated Araport11 database (deposited in the UniProtKB) (Table 1). The results of Hayai-Annotation Plants are presented in Supplementary Data.

**Table 1.** Comparison of annotation of *Arabidopsis thaliana* (Araport11) representative peptide sequences.

|  | Hayai-Annotation<br>Plants<sup>a</sup><br>(4 Best Hits) | Hayai-Annotation<br>Plants<sup>a</sup><br>(20 Best Hits) | Blast2GO<sup>b</sup><br>(Default) | TRAPID |
| --- | --- | --- | --- | --- |
| Specificity (%) | 98.8 | 98.6 | 75.7 | 7.9 |
| Sensitivity (%) | 93.9 | 95.5 | 83.2 | 43.4 |
| Accuracy (%) | 96.4 | 97.1 | 79.4 | 25.7 |
| AED (%) | 3.6 | 2.9 | 20.6 | 74.3 |
| Running time (Total) | 4 m 11 s | 11 m 58 s | 21 h 28 m<sup>*</sup> | 6 h 11 m |
| Running time<br>(Alignment) | 3 m 57 s | 11 m 07 s | 17 h 35 m<sup>*</sup> | NS |
| Running time<br>(Annotation) | 14 s | 51 s | 3 m 53 s | NS |

Note: Mercator 4 only generated EC numbers. FunctionAnnotator were also tested but, after 10 days, retrieved zero results using cDNA file from Araport11 Database. NS: non-specified.
<sup>a</sup> Hayai-Annotation Plants were run on Macintosh laptop (2.2 GHz, 2 cores, 8 GB RAM).
<sup>b</sup> Blast2GO were run on Windows laptop (3.10 GHz, 2 cores, 16 GB RAM). \*BLAST+ was performed on a server (2.40 GHz, 16 cores, 32 GB RAM) with query FASTA file split in 10 (parallel run) in order to increase speed, only for Blast2GO.
<sup>c</sup> TRAPID is a web-based system.

The comparison was performed using all annotated genes, associated with its GO terms, extracted from UniProtKB (regarding Araport11 annotation), against the results obtained from each tested method. Since Hayai-Annotation Plants has the parameter “minimum sequence identity” it was set to 90 % (Blast2GO/BLAST do not have this parameter), with minimum query coverage of 80 %, E-value 1e-6, and “Alignment Score” algorithm. Hayai-Annotation Plants showed a higher accuracy because it selects only the best hit for each GO or EC assignment.

Hayai-Annotation Plants has another major advantage, its speed. The running time for the functional annotation step was four times faster in Hayai-Annotation Plants than Blast2GO. In the sequence alignment step, because USEARCH is already known to be orders of magnitude faster than BLAST (Edgar, 2010), we choose to split the query FASTA file in 10 files to run BLAST+ on a server in a parallel run, the output file, in XML format, was uploaded to Blast2GO.

Furthermore, because the annotation layers are assigned independently, we believe that Hayai-Annotation Plants has the potential to standardize and increase GO terms and EC number, using the “Protein Existence Level” algorithm, inasmuch as minimum sequence identity, minimum query coverage and E-value are outreached.

## Supporting information

## Acknowledgements

We are grateful to Y. Kishida at Kazusa DNA Research Institute for her technical assistance. This work was supported by Life Science Database Integration Project (Database Integration Coordination Program) and Kazusa DNA Research Institute Foundation, Japan.

## Author Contributions Statement

A. G. conceived and S. I. coordinated the project. A. G., K. S., and H. H. analyzed and interpreted the data. A. G. developed the software package. A. G. wrote the manuscript with contributions from K. S., H. H., and S. I. All authors reviewed the manuscript.

