## Supplementary material for "Hayai-Annotation Plants: an ultra-fast and comprehensive gene annotation system in plants": README.pdf

This data set includes nine tables, four graphics, and one data for KEGGmapper.

Table S1 Integrated list of representative genes, from Araport11, with all the annotation layers.

Table S2 A list of Araport11 annotated genes with Gene Ontology Biological Process terms and codes.

Table S3 Count of Gene Ontology Biological Process terms annotated in Araport11.

Table S4 A list of Araport11 annotated genes with Gene Ontology Molecular Function terms and codes.

Table S5 Count of Gene Ontology Molecular Function terms annotated in Araport11.

Table S6 A list of Araport11 annotated genes with Gene Ontology Cellular Component terms and codes.

Table S7 Count of Gene Ontology Cellular Component terms annotated in Araport11.

Table S8 A list of Enzyme Commission annotated in Araport11.

Table S9 Count of Enzyme Commission numbers annotated in Araport11.

Table S10 A data file for KEGG mapper with all EC codes annotated in Araport11.

Graphic S1 Top 50 Gene Ontology Biological Process annotated in Araport11

Graphic S2 Top 50 Gene Ontology Molecular Function annotated in Araport11

Graphic S3 Top 50 Gene Ontology Cellular Component annotated in Araport11

Graphic S4 Top 50 Enzyme Commission annotated in Araport11

EC numbers information is be available in https://enzyme.expasy.org

Supplementary Data.zip

Table S1. hayai_annotation.csv

Table S2. GO_BP_table.csv

Table S3. GO_BP_counts.csv

Table S4. GO_MF_table.csv

Table S5. GO_MF_counts.csv

Table S6. GO_CC_table.csv

Table S7. GO_CC_counts.csv

Table S8. EC_table.csv

Table S9. EC_counts.csv

Table S10. unique_EC.csv

Graphic S1. GO_BP.pdf

Graphic S2. GO_MF.pdf

Graphic S3. GO_CC.pdf

Graphic S4. EC_codes.pdf

README.docx (this file)
