## Supplementary figures and images for "Hayai-Annotation Plants: an ultra-fast and comprehensive gene annotation system in plants"

### EC_codes.pdf

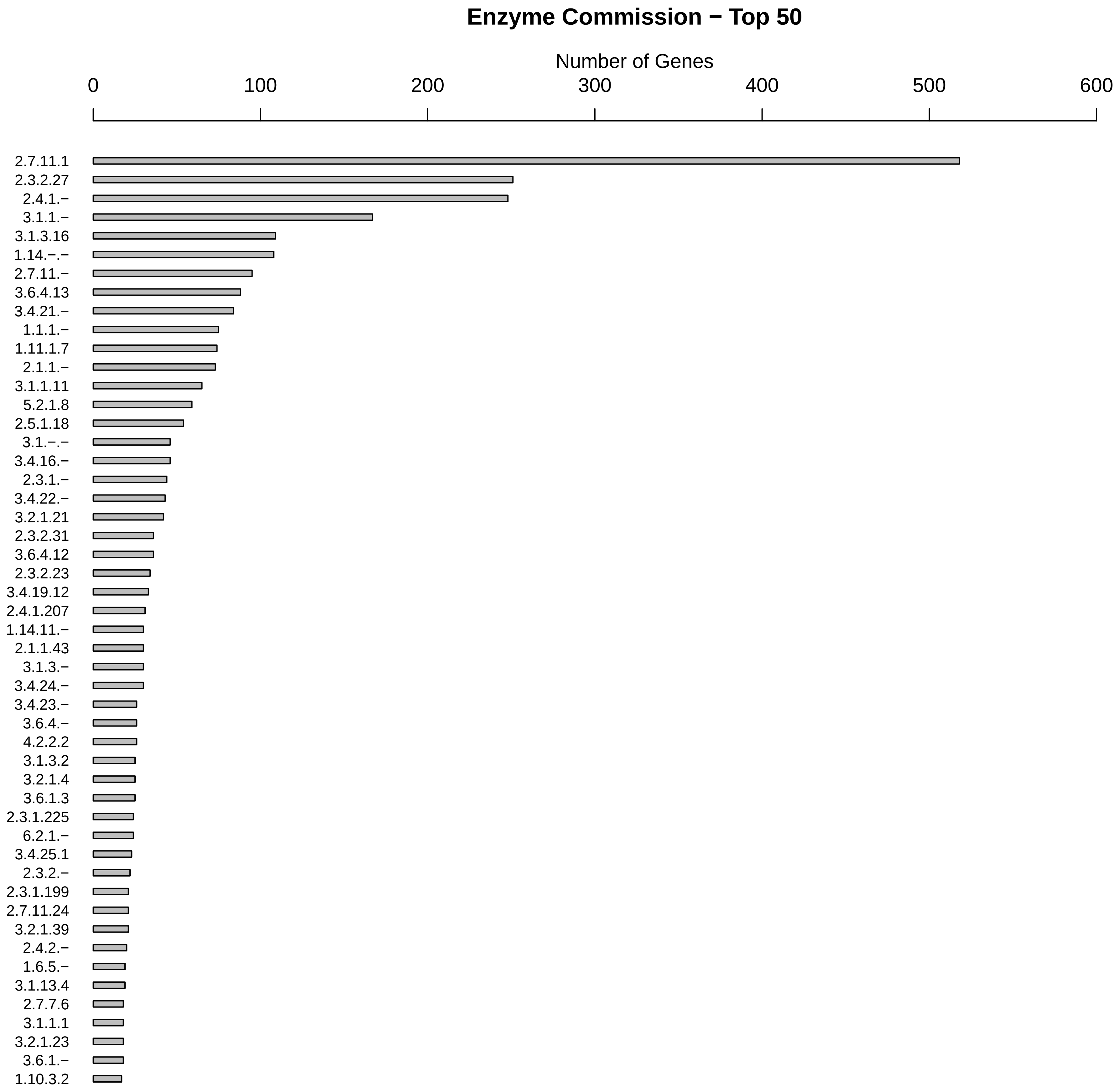

### GO_BP.pdf

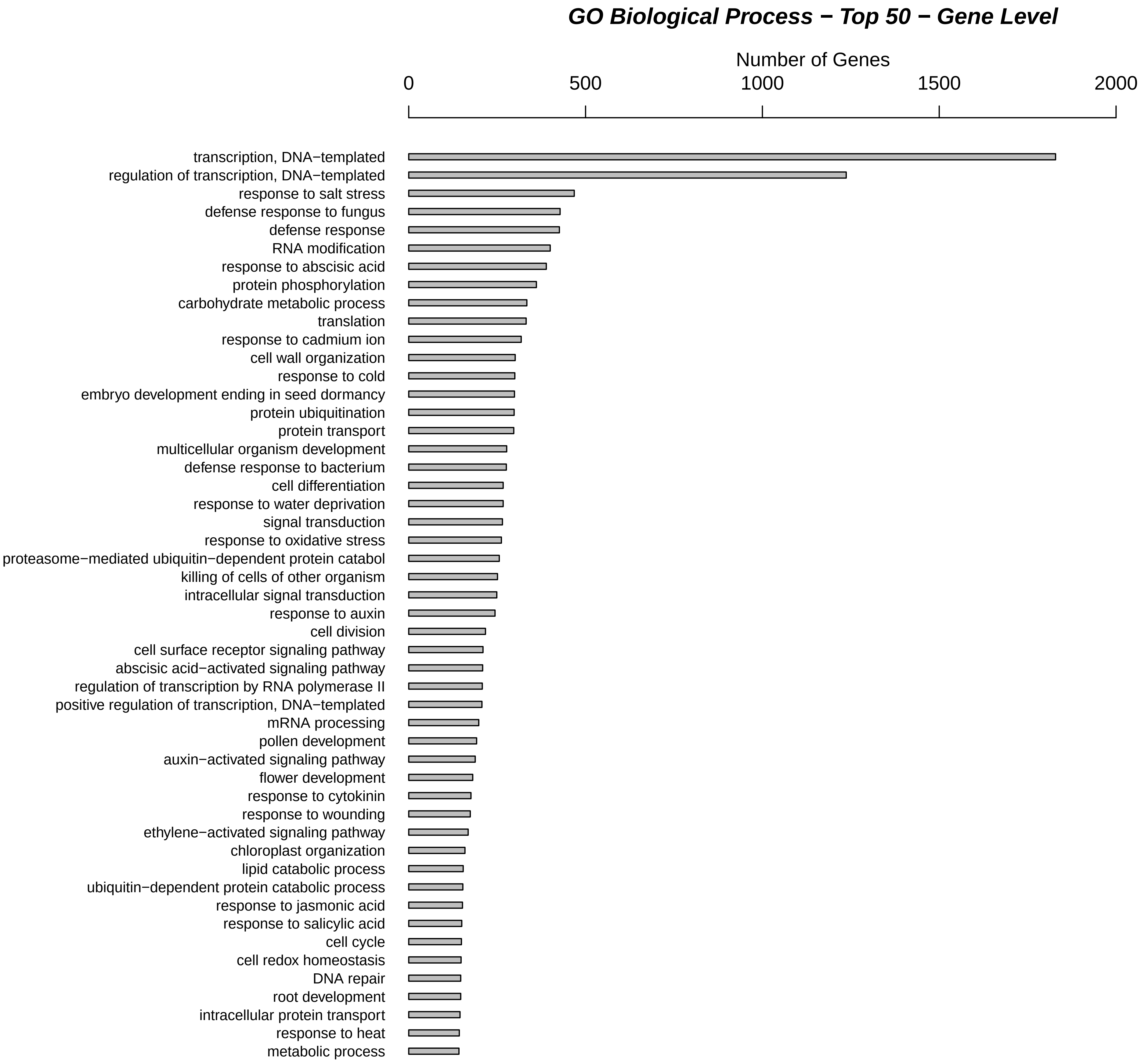

### GO_CC.pdf

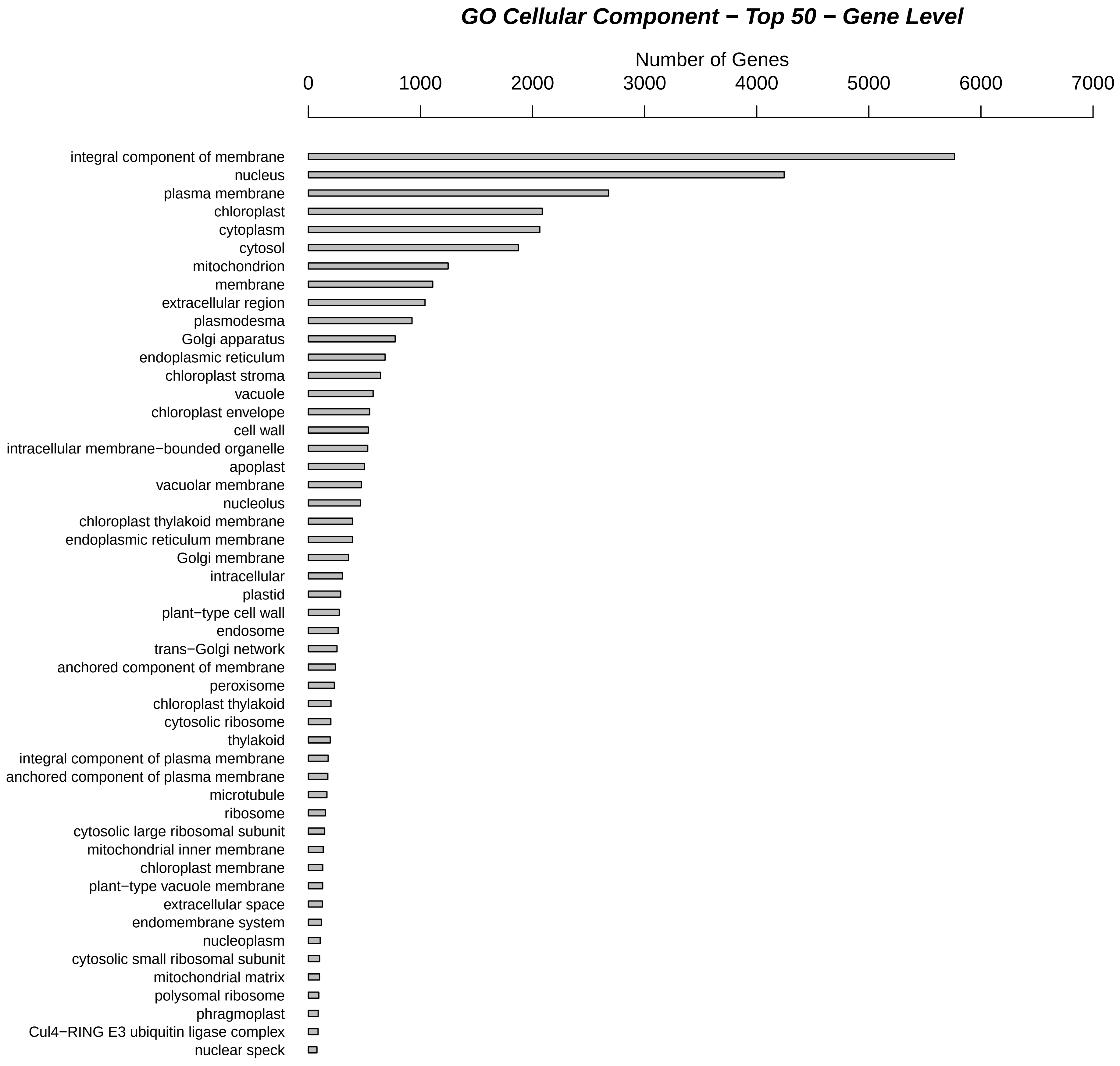

### GO_MF.pdf

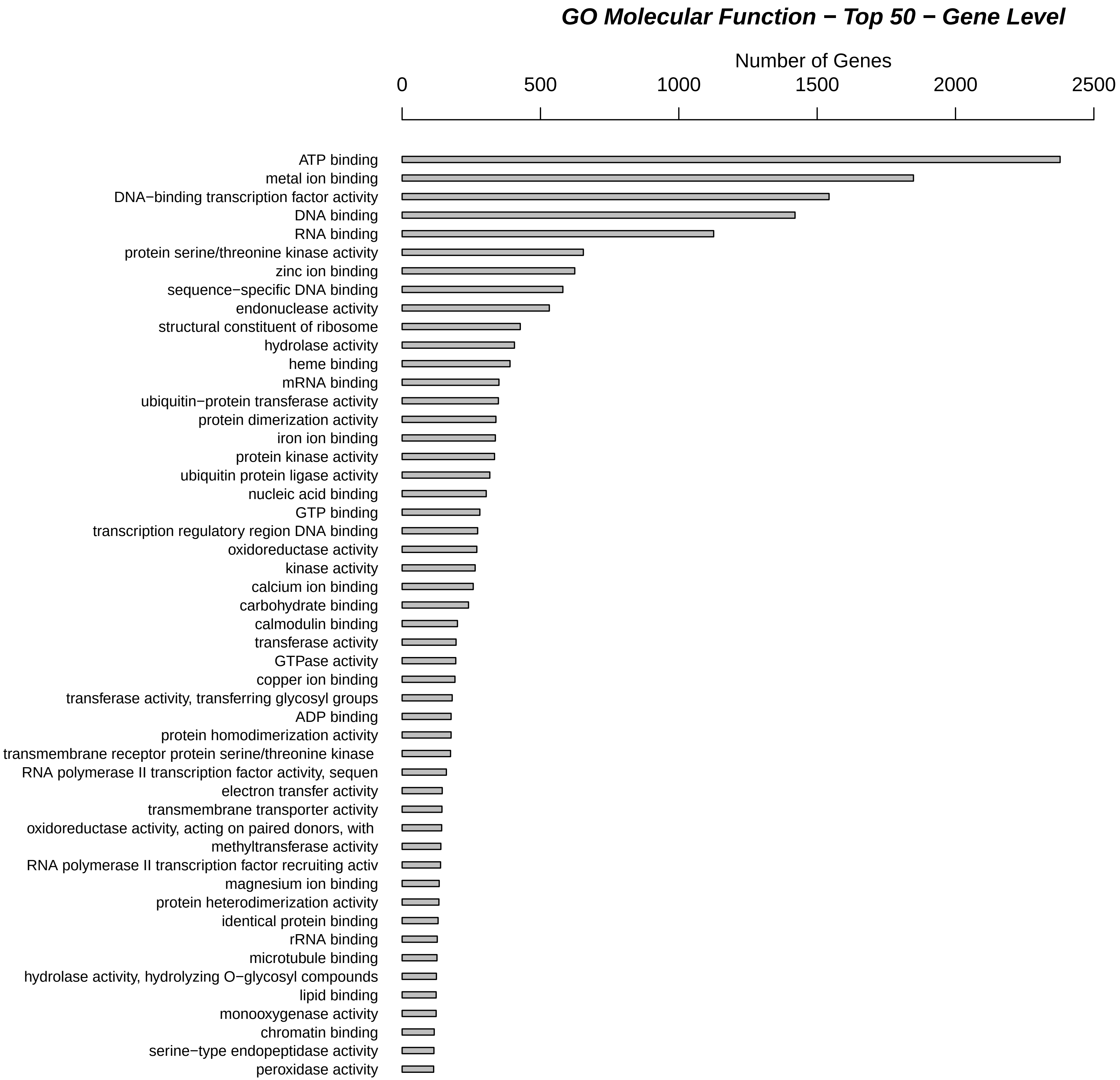
